# Evolution of a Pyroptosis-Apoptosis crosstalk drives non-canonical inflammasome signalling

**DOI:** 10.1101/2025.11.01.685735

**Authors:** Sebastian Grant, Ellis Nicholson, Zoe Hui Xin Chen, Callum Robson, Adam Sung, Barwin Viswanathan, Yunjia Chang, Pranav Patel, James P. Kleppen, Timur Ozdenya, Kaiwen W Chen, Christopher D. Spicer, Dave Boucher

## Abstract

The coordination of efficient immune responses to infections is essential to enable pathogen clearance and host survival. Cell death modalities are a crucial component of the innate immune system and are key to controlling intracellular and extracellular infections. Human caspase-4 and -5 are cysteine proteases activated upon intracellular detection of bacterial lipopolysaccharide (LPS) and triggers pyroptosis, a lytic and pro-inflammatory form of cell death. However, the interaction between these caspases and other cell death modalities remains poorly understood. Here, we build on evidence that caspase-4 can engage in crosstalk with the apoptotic executioner caspases-3 and -7. Our work validated caspase-3 and -7 as direct substrates of caspase-4 and -5, and demonstrated that this specificity was regulated by exosite interactions as opposed to the tetrapeptide. Using AlphaFold, we generated an interaction model between caspase-4 and executioner caspases-3 and -7. From this, we experimentally validated the importance of a hydrophobic interface on caspase-3 and -7 that engaged the caspase-4 exosite, enabling their recognition and cleavage. Importantly, this interface is not used by the apoptotic initiator caspase-8, which cleaved caspase-3 in a tetrapeptide-dependent manner. Our work highlights that inflammatory caspases have evolved a novel mechanism to coordinate crosstalk between pyroptotic and apoptotic signalling, and suggests that these pathways may synergise for the efficient amplification of cell death pathways.

## Introduction

Cell death is critical for multicellular organisms’ homeostasis and defence against insults (1–3). Programmed cell death modalities (PCDMs) have emerged as crucial factors in ensuring homeostasis and responses to infections (2,4). The execution of various PCDMs is controlled by a family of cysteine proteases known as caspases, subdivided into function-associated classes (5,6). Inflammatory caspases are a subgroup of caspases, composed of caspase-1, -4, and -5 in humans, or caspase-11 in mice, that are activated by signalling complexes known as inflammasomes (7–10). Inflammasomes are cellular platforms that sense a range of danger and infection-associated signals and can be classified as either canonical or non-canonical, based on which inflammatory caspases they recruit and activate (9–11). Canonical inflammasomes are composed of host-derived proteins that are activated in response to a plethora of sterile and microbial stimuli, which result in the recruitment and activation of caspase-1. In contrast, the non-canonical inflammasome (NCI) results in the activation of caspases-4, -5 and -11, in response to intracellular lipopolysaccharide (LPS), a cell wall component found on the outer membrane of gram-negative bacteria (12–16). The NCI is particularly critical to immune responses at epithelial barriers during a range of infections in epithelial cells and myeloid cells (17–20). Upon their activation, inflammatory caspases cleave and activate the pro-inflammatory cytokines pro-IL1β and pro-IL18 (13,14,21–25), and the pore-forming protein gasdermin D (GSDMD) (14,26), initiating a highly inflammatory form of cell death known as pyroptosis (27,28). Whilst these are the most well-characterised substrates for inflammatory caspases, proteomic analyses which have established the inflammatory caspase substrate repertoire indicate the presence of multiple signalling axes which can be triggered following their activation (29,30).

The downstream signalling orchestrated by inflammatory caspases ultimately depends on the substrates which are cleaved following their activation. Inflammatory caspase specificity has been shown to be regulated by substrate engagement at two interfaces. All caspases engage their substrates at their substrate-binding pocket, which accommodates four amino acids (P4-P1), known as the tetrapeptide (31). Preference for a specific tetrapeptide sequence varies amongst caspases and has been established using natural and non-natural peptide libraries (32,33). Cleavage sites are generally located in a flexible region, minimising the energy required for catalysis (34). In addition to the tetrapeptide, structural studies have identified other interfaces outside of the active site, known as exosites, which serve as additional determinants mediating the recognition of substrates (24,25,35–37). Interestingly, whilst substrates such as pro-IL18 have been shown to be regulated by both tetrapeptide and exosite interactions (24,25,36), specificity for GSDMD was found to be independent of the tetrapeptide sequence and solely controlled by the exosite (37). The mechanism by which inflammatory caspases recognise other substrates is ill-defined.

In contrast to the inflammatory caspases, which drive pyroptotic cell death, apoptotic caspases classically drive the execution of apoptosis, an immunologically silent form of cell death (4,38). Integrations of apoptotic cues lead to the activation of apical (or initiator) apoptotic caspases-8, -9 and -10 in humans, which cleave the apoptotic executioner caspases-3, -6 and -7, activating them and triggering cellular demise (5). Executioner caspases-3,-6 and -7 require cleavage within their interdomain linker (IDL), which separates their large and small catalytic subunits, to gain activity (5,39). Upon activation, caspase-3 and -7 cleave hundreds of cellular substrates (40,41), including PARP1 (35), ROCK1 (42), the nuclease inhibitor ICAD (43) and the pore-forming protein gasdermin E (GSDME) (44). Whilst caspase-3 is mainly appreciated for its role in apoptosis, cleavage of GSDME by caspase-3 has been shown to drive inflammatory responses via execution of inflammatory PCDMs, such as secondary necrosis (44) and pyroptosis (45).

Despite typically being represented as separate linear pathways, under certain circumstances, caspase-1 has been shown to engage in crosstalk with the apoptotic pathway via both direct and indirect activation of caspase-3 and -7 (46–51). This crosstalk between inflammatory caspases has recently been suggested to extend to the NCI based on recent proteomic data, which flagged caspase-7 as a substrate of caspase-4 (30). However, whether this axis between the NCI, and caspase-3 and -7 is a direct one has not been established. Here, we demonstrate that caspase-4 and -5 directly cleave and activate caspase-3 and -7 at rates comparable to physiologically relevant substrates *in vitro,* and show that the mechanisms underlying this specificity are distinct from caspase-8. Our work provides mechanistic insight into the evolution of a crosstalk between caspase-4 and executioner caspases, and underscores the importance of the roles played by caspase-3 and -7 outside classical apoptotic signalling.

## Methods

### Cell culture

Human monocyte-derived macrophages were produced from PBMC isolated from healthy donors (apheresis product) obtained from the National Health Service. Briefly, PBMCs were separated using a Ficoll layer, washed twice in PBS, and adherent cells were differentiated over 7 days in RPMI 1640 (Gibco) supplemented with 10% foetal calf serum (FCS), 1% glutamine (Gibco), 1% penicillin-streptomycin solution (Gibco), and 50 ng/ml of m-CSF1 (Humankine, Proteintech). HEK293T cells were cultured in DMEM supplemented with 10% FCS, 1% glutamine (Gibco), and 1% penicillin-streptomycin solution (Gibco). Cells were passaged at 70-90% confluency using Trypsin/EDTA (Sigma Aldrich).

### Protein purification

Recombinant caspases were expressed in *E.coli* BL21 as previously described (52). Briefly, a freshly isolated colony was grown in 2YT media overnight and used to seed a larger culture into log phase and induced with 0.2 mM IPTG for 5-18 hrs. After expression, cells were spun at 5000 x g for 5 min, and resuspended in a resuspension buffer (50 mM Tris pH 8.0, 150 mM NaCl). Bacteria were lysed by sonication, purified on a fast-flow Sepharose resin (ThermoFisher), and eluted with imidazole. Protein purity was determined by SDS-PAGE and concentration quantified by spectrophotometry at 280 nm. Active site titration was performed to ensure a robust comparison between the protease batches, as described previously (52). At least two independent batches of caspases were used for most experiments.

### AlphaFold modelling

AlphaFold models were generated using AlphaFold3 (53). Two copies of the human caspase-4 large (a.a 104-270) and small subunit (a.a. 289-377), and the catalytic domains of human caspase-3 or -7 were input for multimer prediction using the default settings. Models which predicted folding of the caspase-4 subunits into the typical caspase heterotetrameric conformation seen in previously published caspase structures were used for further structural analysis in PyMol3 (The PyMOL Molecular Graphics System, Version 3.0 Schrödinger, LLC).

### Immunoblotting

Recombinant proteins or cell extracts were run on an SDS-polyacrylamide gel and transferred to a nitrocellulose membrane using the Trans-Blot Turbo transfer system (Bio-Rad) according to the manufacturer’s instructions. The membranes were blocked in 5% skimmed milk TBST before overnight incubation with the indicated antibodies at 4℃. Antibodies against caspase-4 (anti-mouse, 1:2000, Proteintech), caspase-1 (anti-rabbit, 1:1000, Cell Signaling), GSDMD (anti-rabbit, 1:1000, Cell Signaling), GSDME (anti-rabbit, 1:1000, Proteintech), Histone H3 (anti-mouse, 1:2000, Proteintech), caspase-7 (anti-rabbit, 1:1000, Cell Signaling), caspase-3 (anti-rabbit, 1:1000, Cell Signaling) were used.

### LC-MS

Liquid chromatography-mass spectrometry (LC-MS) was performed on a HCTultra ETD II ion trap spectrometer, coupled to an Ultimate300 HPLC using an Accucore C4 column (100 × 4.6 mm, 5 μm particle size). Water (solvent A) and acetonitrile (solvent B), both containing 0.1% formic acid, were used as the mobile phase at a flow rate of 0.3 mL min^-1^. LC traces were measured via UV absorption at 220, 270, and 280 nm. Spectra were analysed using the Bruker Data Analysis 4.4 software. Spectra were charged deconvoluted using ESI Compass 1.3. Nominal and exact *m/z* values are reported in Daltons.

### Cleavage assays

All cleavage assays were performed in high salt caspase assay buffer (50 mM HEPES pH 7.2,150 mM NaCl, 1 M sodium citrate, 0.01% CHAPS and 10 mM DTT) as described in (52). Briefly, a two-fold dilution series of the indicated concentration of recombinant caspase (as determined by active site titration with Z-VAD-FMK) was performed. Protein substrates were then added with a repeater pipette to a final concentration of 100 nM and incubated for 2 hours at 37°C, except for pro-IL18, which was incubated for 30 minutes. Reactions were terminated by boiling at 95°C for 10 minutes. Samples were then analysed by western blot using the antibodies indicated. Quantification of kinetic parameters was determined by inputting values generated from band intensities using ImageJ, using the following equation to determine an apparent K_cat_/K_M_ as described previously (48).

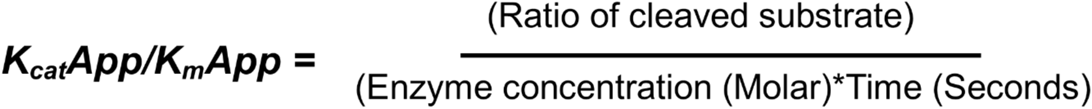

### Zymogen activation assays

Caspase-3 and -7 were expressed as zymogen (unprocessed form) by inducing activation in *E. coli* BL21 for 30 min at 30 °C before purification, as described here (48). Executioner activation was monitored using the caspase-3-like substrate Ac-DEVD-AFC (Sigma-Aldrich) over time, following kinetic reading in a TECAN M200 plate reader. Caspase-4 and zymogen activity on Ac-DEVD-AFC were subtracted from the calculated measurements.

## Results

### Intracellular LPS detection leads to human apoptotic caspase activation

Caspase-1 has been shown to directly cleave caspase-3 and -7 under multiple scenarios (48,49), but this has not been investigated for other inflammatory caspases. The NCI activates caspase-4 and -5 in humans in response to intracellular LPS. To examine if these caspases can also activate caspase-3 and -7, we transfected human monocyte-derived macrophages (HMDMs) with ultrapure *E. coli* K12 LPS. GSDMD cleavage was observed quickly after LPS transfection (Fig1B). Interestingly, we observed the cleavage of caspase-3 and caspase-7 with different kinetics. Caspase-7 was activated at an early time-point (1 hr post-transfection), whereas caspase-3 was cleaved later (3-5 hrs). GSDME, a well-characterised caspase-3 substrate, was also cleaved with similar kinetics as caspase-3 (Fig1A). GSDMD activation downstream of caspase-4 has been shown to activate the NLRP3 inflammasome via GSDMD pore-dependent potassium ion efflux (20). As caspase-1 was shown to directly cleave caspase-3 and -7, we used the NLRP3-selective inhibitor MCC950 (54) to prevent downstream caspase-1 activation. As reported previously (22,55), LPS transfection triggered rapid cell death, which was unaffected by treatment with MCC950, as measured by LDH release (Fig. 1B). In addition, treatment with MCC950 did not prevent GSDME activation following LPS transfection, suggesting that caspase-1 was not critical for the pyroptosis-apoptosis crosstalk observed (Fig1C).

**Figure 1.**
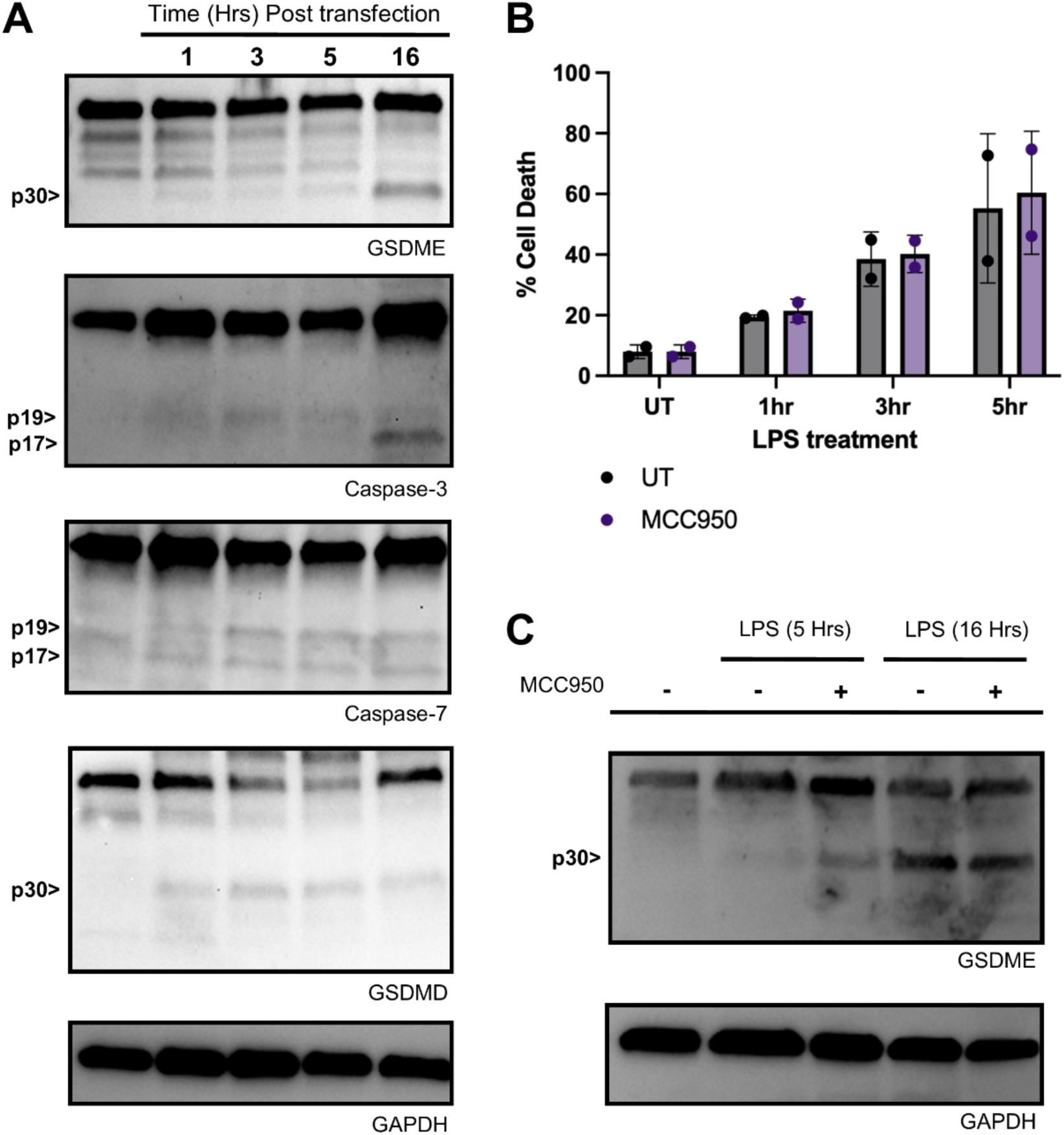
Intracellular LPS detection leads to human apoptotic caspase activation. Human monocyte-derived macrophages were primed with IFNɣ (25 ng/mL, 16hrs) before activation of the non-canonical inflammasome using LPS transfection. A) Kinetic analysis of GSDMD, GSDME, caspase-3 and caspase-7 cleavage over time as assessed by the indicated antibody. B) Lactate Dehydrogenase assay was performed to measure cell death in LPS-transfected macrophages (N=2) in the presence or absence of MCC950 (10 μM) over 5 hours. C) NLRP3 inhibition using MCC950 over 16 hrs does not impact GSDME cleavage as assessed by western blot. Samples were analysed on a 4-15% SDS-PAGE gel. Data is representative of at least three independent experiments from 4-6 individual donors.

**Figure 2.**
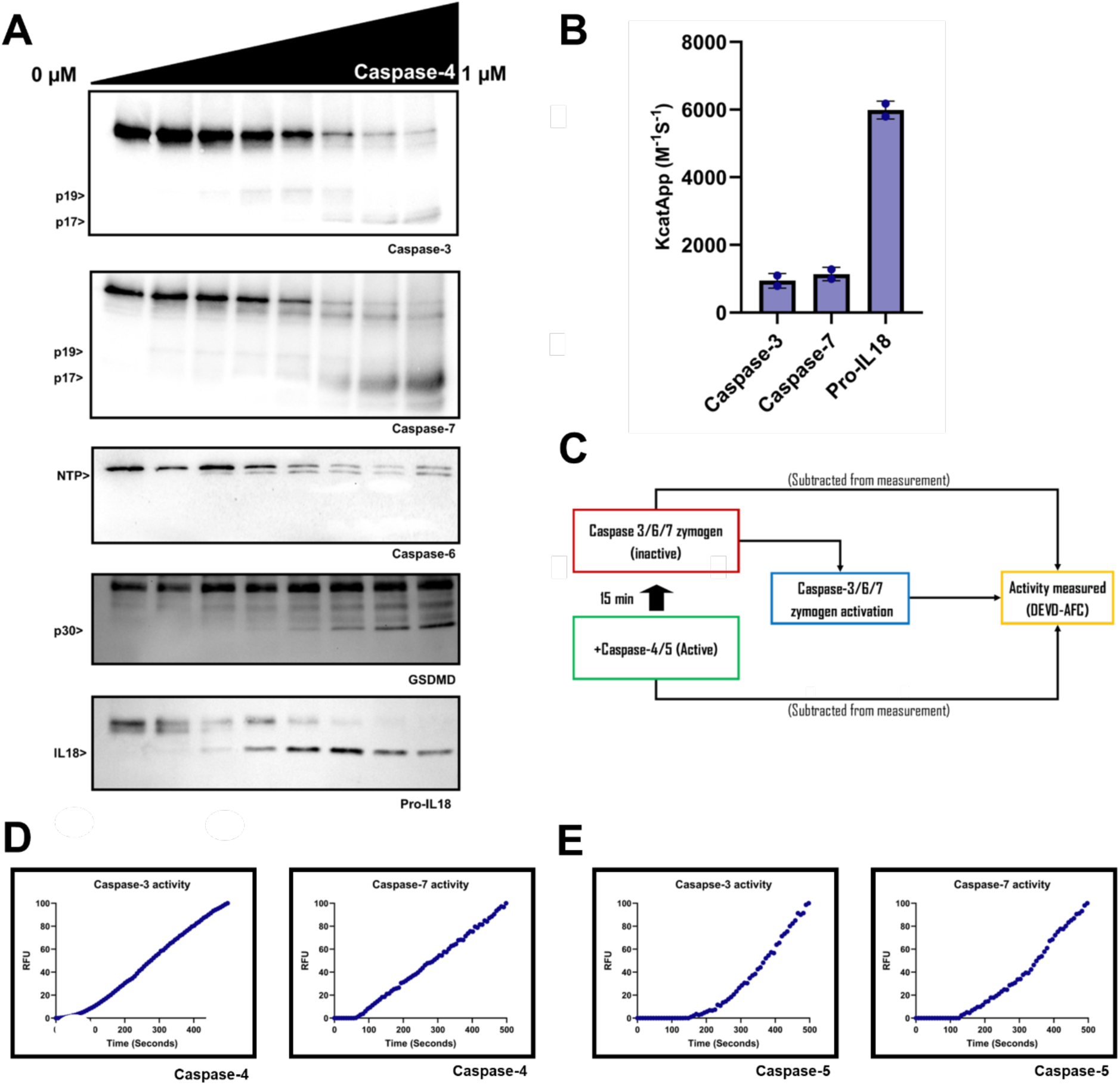
Caspase-4 and -5 directly activate caspase-3 and-7 *in vitro*. **(A)** Western blots of *in vitro* cleavage assays of human caspase-3 (C163A), -6 (C163A), -7 (C186A), GSDMD and pro-IL18, using recombinant caspase-4. **(B)** Quantified processing efficiencies of caspase-3, caspase-7, and pro-IL18 by caspase-4. **(C)** Schematic summary of zymogen activation assays. **(D and E)** Caspase-3 and -7 zymogen activation was measured by relative fluorescence (RFU) generated by Ac-DEVD-AFC substrate cleavage following the assay shown in ‘C’. 1 µM of recombinant caspase-4 **(D)** or -5 **(E)** were used to activate the zymogen preparation. N >= 3 for all experiments.

**Figure 3.**
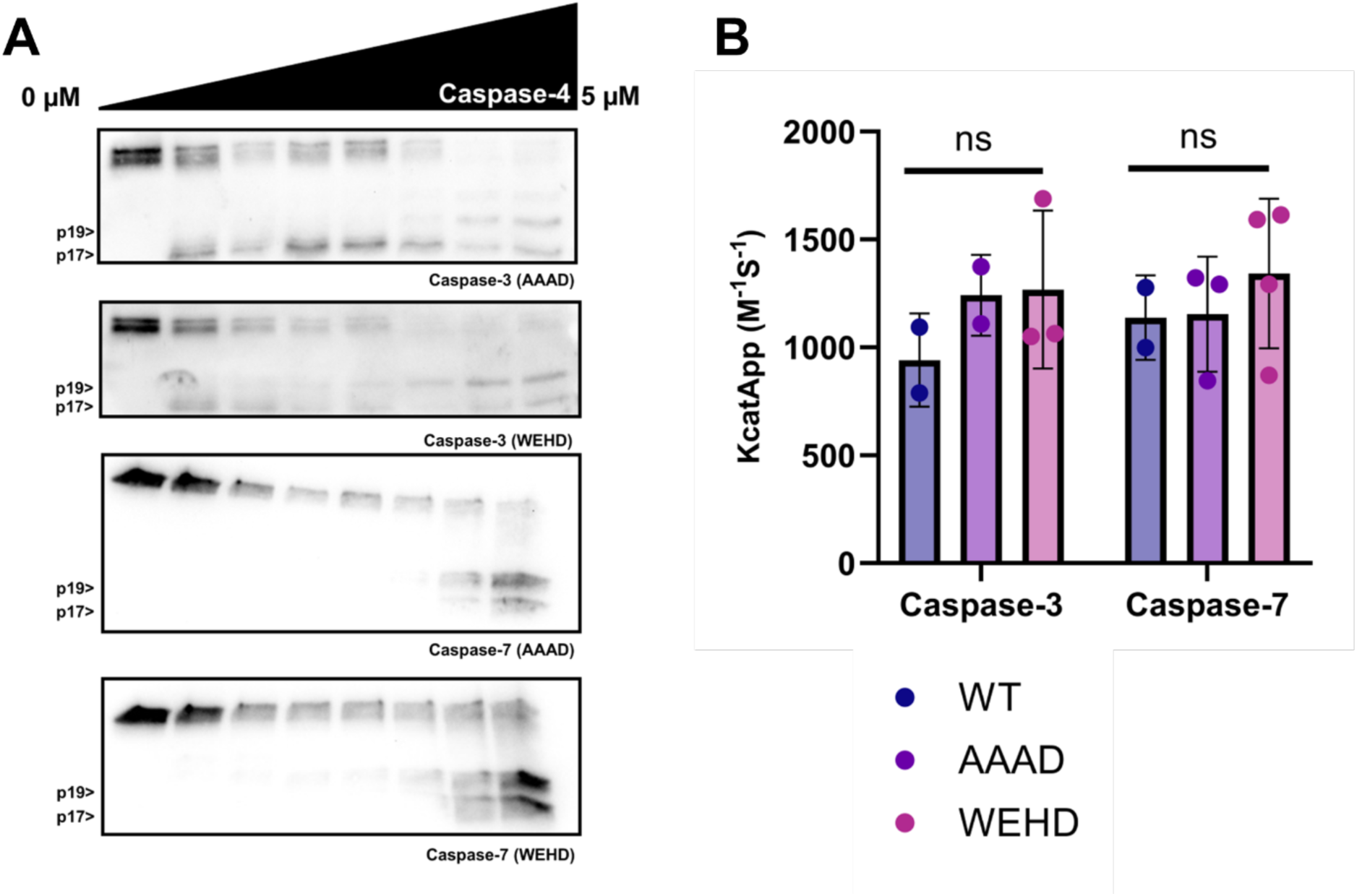
Caspase-4 specificity for caspase-3 and -7 is independent of the tetrapeptide sequence. **(A)** Western blots of in vitro cleavage assays of caspase-3 (C163A) and -7 (C186A) AAAD and WEHD tetrapeptide mutants using recombinant caspase-4. **(B)** Quantified processing efficiencies of caspase-3 and caspase-7 tetrapeptide mutants by caspase-4. Statistical significance assessed by ANOVA p>0.05 (ns). p<0.05 (*), p ≤ 0.01 (**). N=3 at least 3 repeats for each experiments.

**Figure 4.**
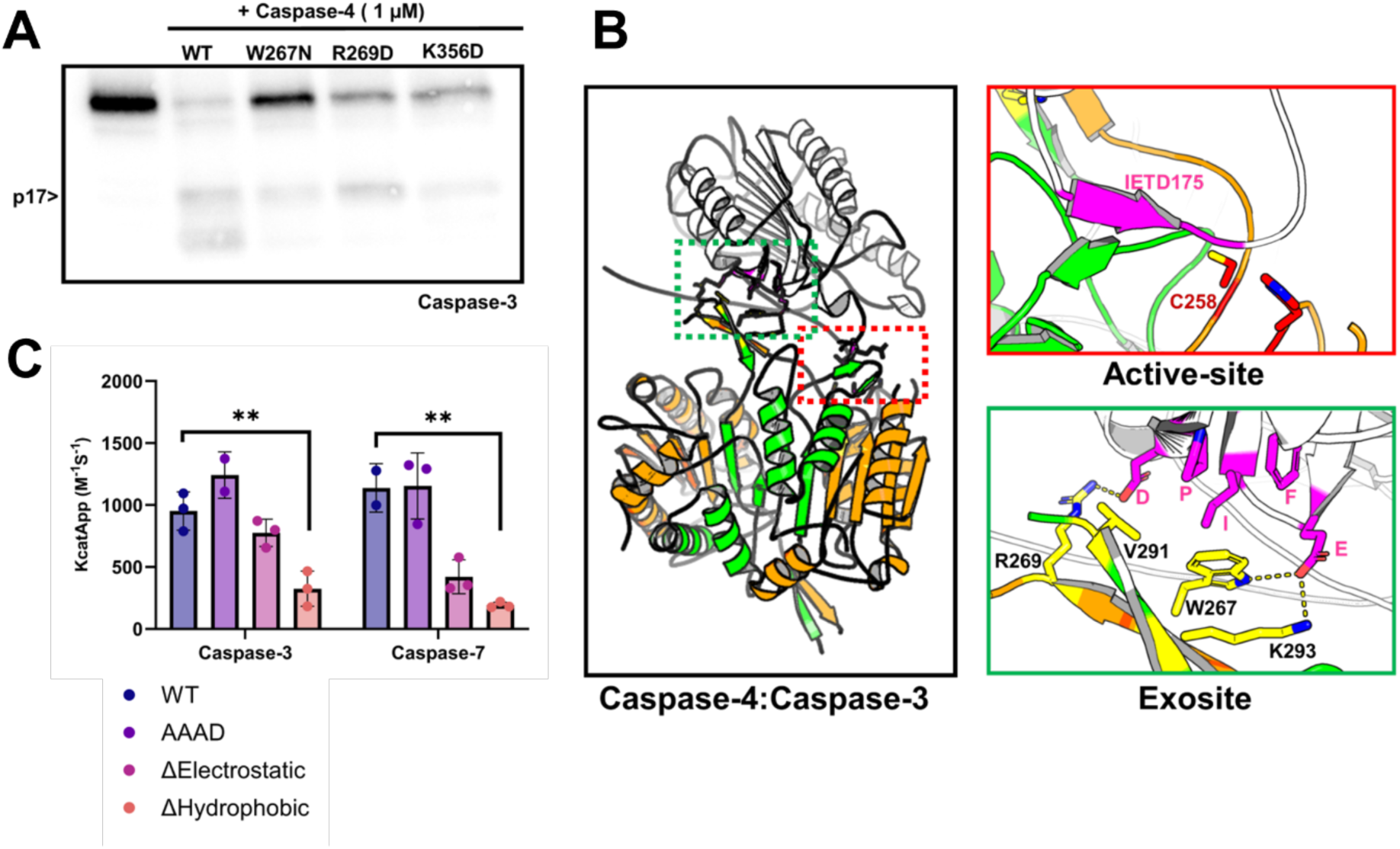
Caspase-4 uses an exosite to confer specificity for caspase-3 and -7. **(A)** Western blot of caspase-3 (C163A) following incubation with 1 µM of recombinant caspase-4 exosite mutants. **(B)** Ribbon representation of AlphaFold multimer model of caspase-4 heterotetramer (orange and green) in complex with caspase-3 (White). The caspase-4 large and small catalytic subunits are highlighted in orange and green, respectively. Red box highlights putative interactions between the caspase-4 catalytic residues (red) and the caspase-3 tetrapeptide (pink). Green box highlights putative interactions at the exosite interface, with caspase-4 residues in yellow, and caspase-3 residues in pink. Figure produced in Pymol3 (Ref). **(C)** Quantified processing efficiencies of caspase-3 and caspase-7 electrostatic and hydrophobic mutants by caspase-4. Statistical significance assessed by ANOVA. p>0.05 (ns). p<0.05 (*), p ≤ 0.01 (**). N >= 3 for all experiments.

### Human caspase-4 and -5 directly activate caspase-3 and -7

Caspase activation kinetics are complex and can result from multiple signalling events. For example, non-caspase proteases (e.g., Granzyme B (56)) have been shown to activate caspase-3 and -7 directly. To confirm that caspase-4 can directly cleave caspase-3 and -7, we incubated recombinant catalytically inactive caspase-3 (C163A), -6 (C163A) and -7 (C186A) with recombinant caspase-4 for 2 hrs (Fig2A). Executioner caspases have been shown to be cleaved at defined aspartate residues, leading to their proteolytic activation (5). Upon incubation, we observed the generation of 19 and 17 kDa bands for caspase-3 and -7 by western blot, indicating potential activation, similar to the pattern created by caspase-8 and caspase-9 cleavage (5,39). However, contrary to caspase-8 and -9, caspase-4 could also cleave the N-terminal peptide of caspase-7, generating the fully active form (39). During apoptosis, such fragments are normally cleaved by caspase-3 to generate a fully active caspase-7. We also observed cleavage of caspase-6 following caspase-4 incubation; however this did not result in the cleavage of its inter-domain linker (IDL) but only of its N-terminal peptide. To compare how efficient caspase-3 and -7 were cleaved compared to other notable substrates of caspase-4, we calculated kinetic parameters for processing of caspase-3 and -7 (Fig2B). Upon quantification of processing efficiencies, caspase-3 and-7 were found to be processed slower than pro-IL18, but more efficiently than other reported substrates such as pro-IL-1β and pro-IL-36y (24). We confirmed the location of caspase-4 cleavage sites on caspase-3, -6 and -7 by liquid-chromatography mass spectrometry (LC-MS). Molecular masses which corresponded to cleavage at the canonical IDL activation sites for caspase-3 (D175) and -7 (D198) were detected (FigS1). LC-MS confirmed that caspase-4 cleaved D23 within the N-terminal peptide of caspase-6 *in vitro*. To confirm whether cleavage of caspase-3, -6 and -7 by caspase-4 led to their proteolytic activation, we performed zymogen activation assays in which enzyme activity of zymogen preparations were measured with and without prior incubation with caspase-4 (Fig2C) (52). Proteolytic activation of caspase-3 and -7 was confirmed by an increase in caspase-3 and -7 protease activity when their zymogens were incubated with recombinant caspase-4 (Fig2D). Similar processing and activation of caspase-3 and -7 was observed following incubation with caspase-5 (Fig2E, SupFig2). Together, these data demonstrate that caspase-3 and -7 are direct substrates of caspase-4 and -5 and are processed at rates comparable to other physiologically relevant substrates *in vitro*.

### Caspase-4 uses an exosite to confer caspase-3 and -7 specificity

Inflammatory caspases have been shown to utilise a binary mechanism for conferring substrate specificity. Both GSDMD and pro-IL18 have been shown previously to utilise an exosite to mediate their recognition by inflammatory caspases with differential dependencies on the tetrapeptide sequence (24,36,37). Pro-IL18 specificity is regulated by both exosite and tetrapeptide interactions, whereas GSDMD processing appears to be completely independent of the tetrapeptide sequence and is entirely controlled by the exosite. To establish the mechanistic details underpinning caspase-4’s specificity for caspase-3 and -7, we generated mutants of caspase-3 and -7 in which the tetrapeptide was mutated to either an optimal WEHD or suboptimal AAAD sequence. Similarly to what has been reported for GSDMD, mutation of the caspase-3 and -7 tetrapeptide to AAAD did not affect their cleavage by caspase-4 (Fig3A). Additionally, upon calculation of kinetic parameters, we observed no significant difference in caspase-3 or -7 processing between the WT, WEHD and AAAD tetrapeptide mutants, indicating that specificity for caspase-3 and -7 was not regulated by the tetrapeptide sequence (Fig3B).

This prompted us to analyse whether the caspase-4 exosite may be conferring substrate specificity. Residues W267, R269 and K356 are residues within the caspase-4 exosite which were previously found to mediate specificity for pro-IL18 and GSDMD (24,36,37). We expressed caspase-4 mutants carrying charge or polarity switch mutations in these exosite residues (W267N, R269D, K356D). When compared, mutation of W267 to a polar residue resulted in the largest reduction in caspase-3 processing, with charge-reversion of K356 and R269 having little effect on processing (Fig4A). These findings suggest that W267 is the primary residue conferring specificity for caspase-3, with K356 and R269 being less critical. To understand how W267 engaged caspase-3 to confer specificity, we generated AlphaFold models of the caspase-4 heterotetramer in complex with caspase-3 and -7 (Fig4B, SupFig3). Consistent with our experimental findings, the models successfully predicted engagement of the caspase-3 and -7 IDL, which contained the D175 and D198 cleavage site within the caspase-4 active site, respectively. Upon inspection of the caspase-4 exosite, all of the generated models mapped W267 in proximity to hydrophobic residues present on caspase-3 composed of I126, F128, and P133. When we mutated these residues that were predicted to engage W267 (I126G, F128G, P133G, termed caspase-3/7 ‘hydrophobic mutant’), we observed a significant reduction in caspase-4 processing of both caspase-3 and -7, indicating that these residues were indeed important for specificity (Fig4C). The models also mapped R269 and K320 in proximity to negatively charged residues (E123 and D135, respectively), which could engage in electrostatic interactions as seen in the caspase-4:pro-IL18 structures (24,36). However, in accordance with our experimental observations that R269 was less important for specificity, we did not observe significant reductions in processing when charge-reversing mutations were made at this interface on caspase-3 (E123R, D135R) or -7 (E146R, D161R) (Fig4C). Together, these data suggest that the caspase-4 uses W267 within its exosite to confer specificity for caspases-3 and -7 via engagement of hydrophobic residues.

### Caspase-8 uses the tetrapeptide to confer specificity for caspase-3 and GSDMD

Following confirmation that caspase-4 used an exosite to recognise caspase-3 and -7, we wanted to test whether caspase-8 used similar mechanisms to confer specificity for caspase-3. To do this, we performed cleavage assays using recombinant caspase-8 on our caspase-3 mutants as substrates. Surprisingly, unlike caspase-4, mutation of the caspase-3 tetrapeptide to AAAD abrogated caspase-8 cleavage of caspase-3, indicating that its specificity was dependent on the tetrapeptide (Fig5A). Additionally, the caspase-3 hydrophobic mutant, which reduced processing by caspase-4, was cleaved by caspase-8 to a similar extent as wild-type caspase-3, indicating that caspase-8 did not use this interface to mediate recognition (Fig5B). This difference in substrate recognition of redundant substrates also extended to GSDMD, which has been previously shown to be cleaved by both caspase-8 and inflammatory caspases (57,58). Once again, unlike the inflammatory caspases, which cleaved GSDMD in a tetrapeptide-independent manner, mutation of the GSDMD tetrapeptide to AAAD reduced its cleavage by caspase-8 (SupFig4). Finally, to speculate whether this axis between the NCI and caspase-3 may be conserved, we assessed the conservation of residues at the caspase-4:caspase-3 exosite interface across mammals. We found strong conservation of W267, consistent with it having central roles for GSDMD, pro-IL18, and caspase-3 specificity. Notably, R269 was not strongly conserved across the mammalian lineage, indicating that its contribution at the exosite may be limited to certain signalling axes such as pro-IL18 processing (Fig. 5C). When assessing conservation of residues on caspase-3 which were found to be important for its processing by caspase-4, we found conservation of hydrophobic residues across the mammalian lineage, indicating that the caspase-4:caspase-3 signalling axis may also be present within other mammalian species (Fig5C). Together, these results reveal molecular and evolutionary insight into caspase-4-dependent crosstalk with apoptotic caspases-3 and -7 and discern the mechanisms of caspase-4’s specificity for caspase-3 and -7, which are distinct from caspase-8.

**Figure 5.**
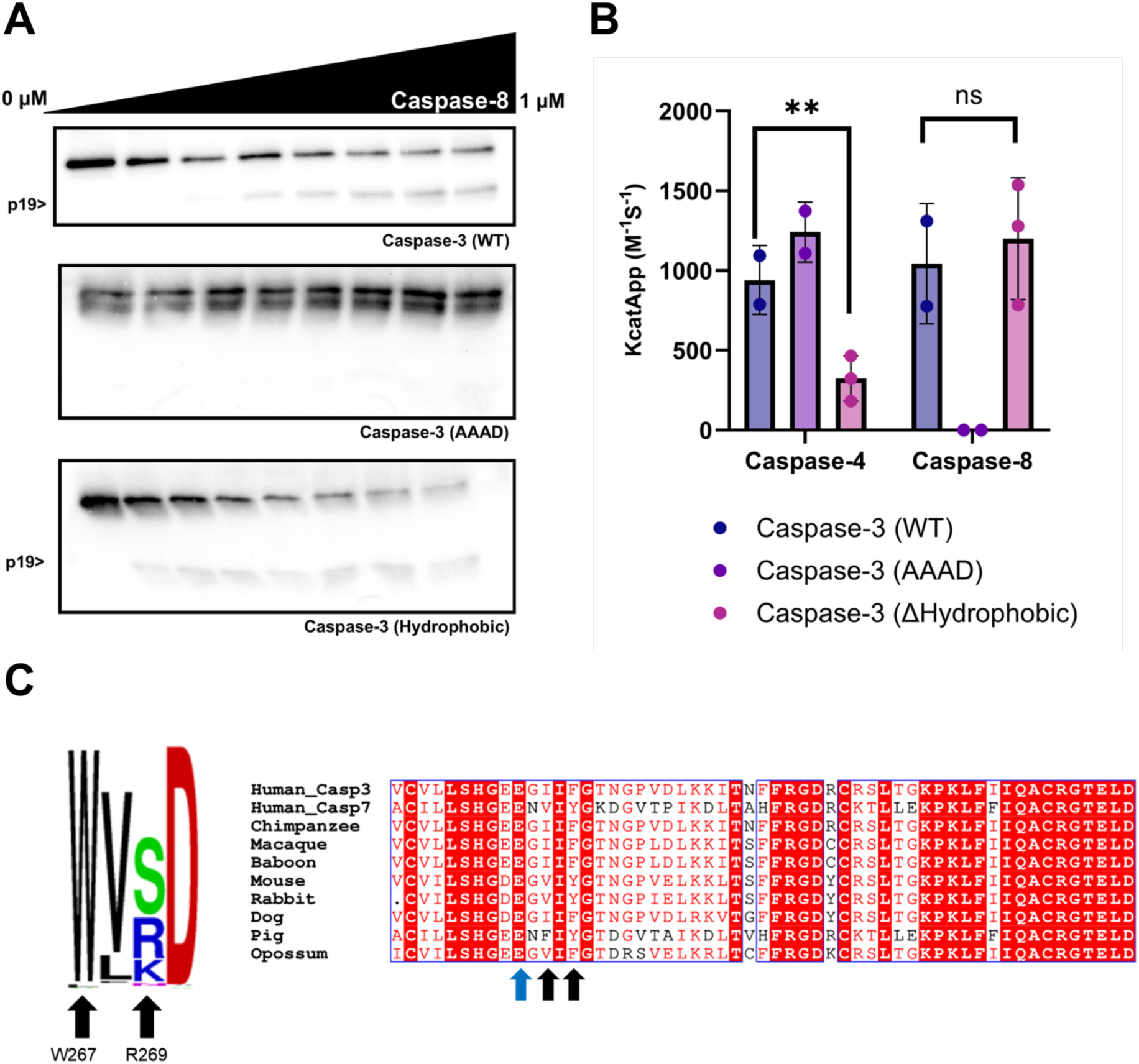
Caspase-8 uses distinct mechanisms to confer caspase-3 specificity. **(A)** Western blots of *in vitro* cleavage assays of caspase-3 (C163A) AAAD and hydrophobic mutants using recombinant caspase-8. **(B)** Quantified processing efficiencies of caspase-3 mutants by caspase-4 and -8. Statistical significance assessed by ANOVA. p>0.05 (ns). p<0.05 (*), p ≤ 0.01 (**). **(C)** (Left) Residue conservation of the caspase-4 exosite across mammals, with letter height indicating relative frequency at each position. (Right) Sequence alignment demonstrating conservation of exosite-engaging residues (indicated by black arrows for W267-engaging and blue arrive for R269-engaging) across mammalian caspase-3 homologous. N >= 3 for all experiments.

## Discussion

The substrate specificity of caspases ultimately determines their downstream function upon their activation. Caspase-4 activation during Gram-negative bacterial infection triggers a signalling axis leading to the activation of apoptotic caspases-3 and -7, independent of caspase-1. However, the mechanisms underpinning this specificity for caspase-3 and -7, and the efficiencies at which they are processed had not previously been explored. Here, we demonstrated that caspase-3 and -7 were direct substrates of caspase-4 and -5, and their processing resulted in their activation. We calculated the catalytic parameters for this processing which enabled comparison to other known substrates of caspase-4 and -5, demonstrating that caspase-3 and -7 are processed at rates comparable to other known physiologically relevant substrates rates. Interestingly, although caspase-3 and -7 appeared to be processed to similar efficiencies *in vitro*, we observed more rapid activation of caspase-7 following LPS transfection in cells, indicating that factors outside of substrate specificity likely influence activation kinetics in cellular contexts. Mechanisms which modulate substrate localisation, abundance, and phosphorylation, are all factors which may be involved in such signalling biases (59). The role of caspase-3 synergism with the pyroptotic pathway via GSDME activation is well understood, providing a clear evolutionary rationale for its activation by caspase-4 during bacterial infection. However, despite caspase-7 having the capacity to activate GSDME in most vertebrates, it has been reported to have lost this ability in the primate lineage due to acquisition of mutations bordering its substrate binding pocket (60,61). This makes the downstream phenotypic consequences of caspase-7 activation by caspase-4 in primates less clear. The cleavage of Acid Sphingomyelinase (62) and other caspase-7 specific substrates may help balance inflammation states, however all these downstream consequences warrant additional investigation.

We demonstrated that, in contrast to caspase-8, caspase-4 did not use the tetrapeptide to enable recognition of caspase-3 and -7, and instead used the same exosite which mediates recognition of GSDMD and pro-IL18 (24,36,37). Point mutations within the caspase-4 exosite identified W267 as the primary residue contributing to caspase-3 specificity, with K356 and R269 having less prominent contributions. The molecular landscape predicted in the generated three-dimensional models is reminiscent of experimentally established structures of caspase-4 with pro-IL18 and GSDMD, with W267 engaging hydrophobic regions on the substrate (24,36,37). Additionally, despite appearing to be less important for caspase-3 processing, putative electrostatic interactions with R269 and K320, which flank the hydrophobic patch of caspase-3 were predicted in the models, similar to those interactions seen within the pro-IL18:caspase-4 crystal structures (24,36). Whilst the exact molecular landscape cannot be accurately predicted without experimental models, it can be speculated that the hydrophobic region engaged by W267 likely arises from I126 and the phenyl of F128 in caspase-3. F128 is substituted for a tyrosine in caspase-7 and some mammalian caspase-3 homologues, suggesting that the hydrophobic phenyl ring, which is common to both tyrosine and phenylalanine, participates in exosite engagement. Additionally, the preference for an isoleucine does not appear to be exclusive, with substitutions for other hydrophobic residues (valine in caspase-7) being present in caspase-7 and caspase-3 orthologues. Indeed, whilst these residues were found to be important for specificity, it was noted that mutation of this interface alone did not abolish cleavage by caspase-4, indicating that other interactions at the caspase-4 exosite are likely contributing towards specificity.

The reason why caspase-4 has evolved a distinct mechanism from caspase-8 to target caspase-3 is unclear, however it is likely due to activation of the caspase-3/GSDME axis during bacterial infection being important for fitness (2). This is supported by the conservation of the caspase-3/GSDME axis across the vertebrates (60). Indeed, it can be speculated that alteration to the inflammatory caspase substrate binding pocket to encompass new functions such as IL-18 maturation may correlate with a loss in ability to activate the GSDME axis via tetrapeptide-mediated caspase-3 activation. Accordingly, the evolution of an exosite circumvents this trade-off by enabling the acquisition of caspase-3 and GSDMD processing capabilities, whilst maintaining its new primary function of IL-18 maturation. This trade-off is likely crucial in controlling infections by Gram-negative bacteria, as recently demonstrated for *Salmonella* (*63*). This is consistent with the hypothesis that the evolution of cytokine processing abilities must have co-evolved with a mechanism to enhance the export of mature cytokines, such as simultaneous activation of gasdermin pores upon inflammatory caspase activation (61).

Together, our findings reveal mechanistic insights into how inflammatory caspases orchestrate crosstalk with caspase-3 and -7 and show that the mechanisms employed are distinct from other initiator caspases, such as caspase-8. Additionally, this substrate-specific distinction provides evolutionary insight into why inflammatory caspases may have evolved an exosite to target specific substrates.

## Contribution

SG, ZC, EN, CR, BV, PP, JK, YC, TO, and DB performed experiments and analysed data. CS and KWC supervised part of the project and provided essential feedback and reagents. DB designed and supervised the project and secured funding. DB and SG wrote the manuscript with input from all contributors. The authors have no conflict of interest to declare.

## Acknowledgements

This work was supported by a BBSRC project grant (BB/Y009703/1) and a Royal Society project grant to DB (RGS\R1\231411). SG is supported by a doctoral training scholarship from the York Graduate Research School and the Yorkshire Consortium for Equity in Doctoral Education (YCEDE). BV and JPK are supported by WhiteRose BBSRC PhD studentships. AS and PP were supported by a YCEDE summer studentship. Chris and Kaiwen to add their funding. CDS acknowledges generous support through a Wellcome Trust Career Development Award (225257/Z/22/Z). KWC is funded by a MOE Tier 1 grant (NUHSRO/2025/011/T1/Seed-Sep24/01). ZHXC is supported by a NUSMed Research Scholarship. We thank Dr Ed Bergstrom and The York Centre of Excellence in Mass Spectrometry for support with protein mass spectrometry. This centre was created thanks to a major capital investment through Science City York, supported by Yorkshire Forward with funds from the Northern Way Initiative, and subsequent support from the EPSRC (EP/K039660/1; EP/M028127/1). The authors thank the Taabazuying lab for helpful discussions and comments on the manuscript. The authors thank members of the Boucher lab, Spicer lab, Dr Jillian Barlow and Dr Kim Robinson for helpful discussions.

---

**Supplementary figures.**
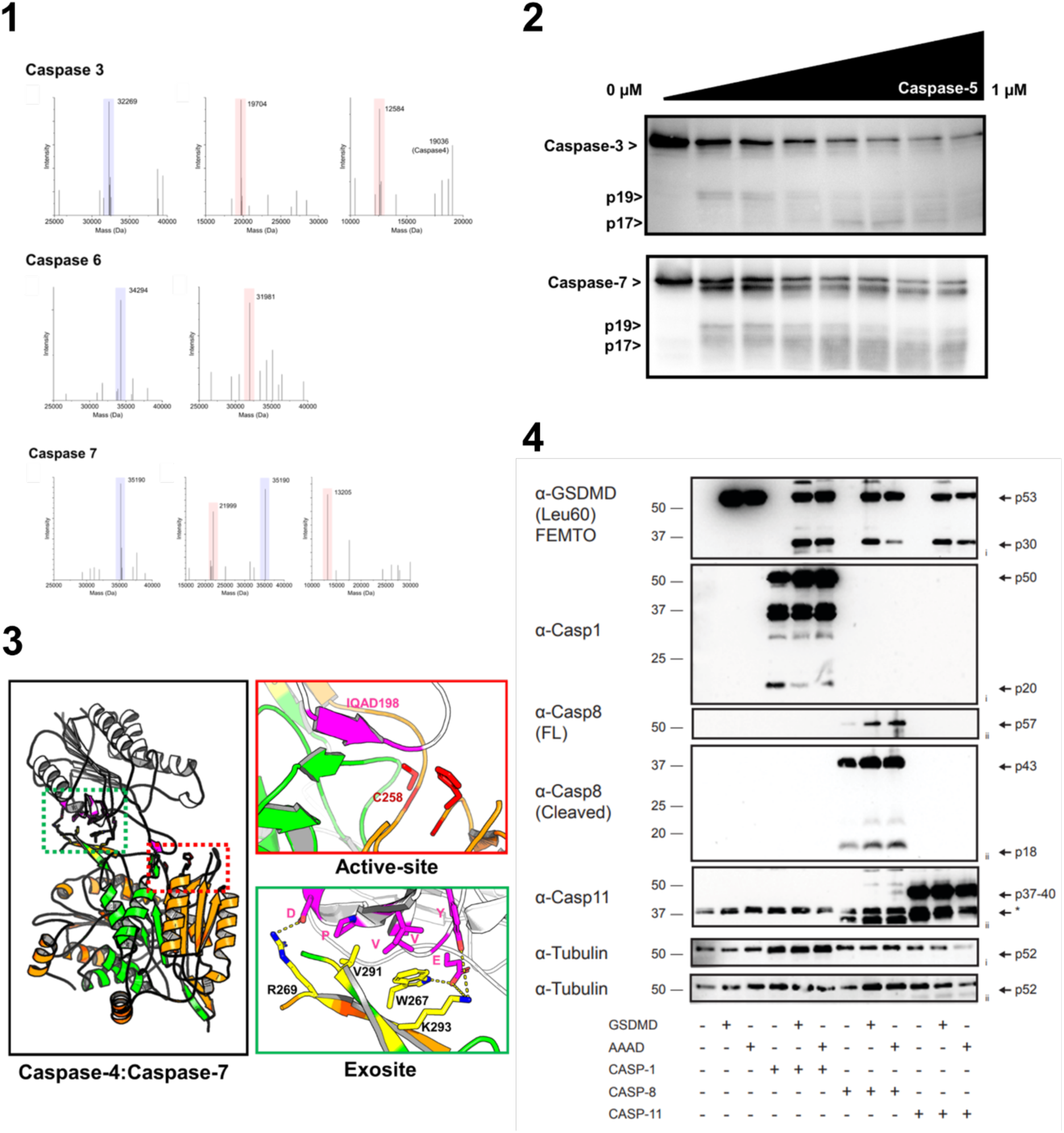
**(1)** Deconvoluted mass spectra derived from LC-MS analysis of peptide products generated from reactions containing human caspase-3 (C163A), -6 (C163A), -7 (C186A) with and without incubation with recombinant caspase-4. Peaks highlighted in blue indicate the undigested substrate, whilst peaks highlighted in red indicate cleavage products generated following incubation with caspase-4. **(2)** Western blots of *in vitro* cleavage assays of caspase-3 (C163A) and -7 (C186A), mutants using recombinant caspase-5. **(3)** Ribbon representation of AlphaFold multimer model of caspase-4 heterotetramer (orange and green) in complex with caspase-7 (White). The caspase-4 large and small catalytic subunits are highlighted in orange and green, respectively. Red box highlights putative interactions between the caspase-4 catalytic residues (red) and the caspase-7 tetrapeptide (pink). Green box highlights putative interactions at the exosite interface, with caspase-4 residues in yellow, and caspase-7 residues in pink. Figure produced in Pymol3. **(4)** HEK 293T cells co-expressing GSDMD WT or an AAAD tetrapeptide mutant of GSDMD, and ΔCARD DmrB-Caspase-1, -8 or -11. Cells were then treated with 1 µM AP20187 to activate the DmrB-caspases for 1h and lysates were assessed by western blot.

